# Reduced representation sequencing for symbiotic anthozoans: are reference genomes necessary to eliminate endosymbiont contamination and make robust phylogeographic inference?

**DOI:** 10.1101/440289

**Authors:** Benjamin M. Titus, Marymegan Daly

## Abstract

Anthozoan cnidarians form the backbone of coral reefs. Their success relies on endosymbiosis with photosynthetic dinoflagellates in the family Symbiodiniaceae. Photosymbionts represent a hurdle for researchers using population genomic techniques to study these highly imperiled and ecologically critical species because sequencing datasets harbor unknown mixtures of anthozoan and photosymbiont loci. Here we use range-wide sampling and a double-digest restriction-site associated DNA sequencing (ddRADseq) of the sea anemone *Bartholomea annulata* to explore how symbiont loci impact the interpretation of phylogeographic patterns and population genetic parameters. We use the genome of the closely related *Exaiptasia diaphana* (previously *Aiptasia pallida)* to create an anthozoan-only dataset from a genomic dataset containing both *B. annulata* and its symbiodiniacean symbionts and then compare this to the raw, holobiont dataset. For each, we investigate spatial patterns of genetic diversity and use coalescent model-based approaches to estimate demographic history and population parameters. The Florida Straits are the only phylogeographic break we recover for *B. annulata*, with divergence estimated during the last glacial maximum. Because *B. annulata* hosts multiple members of Symbiodiniaceae, we hypothesize that, under moderate missing data thresholds, *de novo* clustering algorithms that identify orthologs across datasets will have difficulty identifying shared non-coding loci from the photosymbionts. We infer that, for anthozoans hosting diverse members of Symbiodinaceae, clustering algorithms act as *de facto* filters of symbiont loci. Thus, while at least some photosymbiont loci remain, these are swamped by orders of magnitude greater numbers of anthozoan loci and thus represent genetic “noise,” rather than contributing genetic signal.

## 1. Introduction

The study of the distribution of genetic diversity across broad geographic space (i.e. phylogeography) can shed light on the historical and contemporary processes responsible for the generation and maintenance of biodiversity within species and ecosystems (Arbogast, 2001; Avise, 2009; Avise, Bowen, & Ayala, 2016; Knowles, 2009). Phylogeographic surveys demarcate barriers to dispersal, routinely recover cryptic species, and with increasing dataset sizes and statistical approaches, can estimate important demographic parameters such as effective population size, divergence time, migration rates, and historical changes in population size (e.g. Avise et al., 2016; Carstens, Pelletier, Reid, & Satler, 2013; Knowles, 2009; Pante et al., 2015; Pelletier & Carstens, 2014; Smith et al., 2017). To that end, high-throughput sequencing, which can generate thousands of unlinked single nucleotide polymorphisms (SNPs) across the genome, has been particularly powerful, allowing for greater statistical and explanatory power into complex evolutionary and demographic histories (e.g. Carstens, Lemmon, & Lemmon, 2012; Excoffier, Doupenloup, Huerta-Sánchez, Sousa, & Foll, 2013; McCormack, Hird, Zellmer, Carstens, & Brumfield, 2013).

Although the field of phylogeography has a long history in marine systems (e.g. Bowen et al., 1992, 1994; Reeb & Avise, 1990), cnidarians in the class Anthozoa (i.e. corals, sea anemones, zoanthids, corallimorpharians), which form the backbone of coral reefs and a major component of its biodiversity, have been historically challenging to work with at the population level. In addition to large range sizes and the logistical difficulties of sampling underwater, mitochondrial DNA barcodes (mtDNA), the molecular marker of choice for metazoan phylogeographic studies from the field’s outset, evolve too slowly in most anthozoans to be useful for intraspecific studies (e.g. Allio, Donega, Galtier, & Nabholz, 2017; Daly, Gusmão, Reft, & Rodríguez, 2010; Shearer, Van Oppen, Romano, & Wörheide, 2002;). Further, the overwhelming majority of tropical anthozoans found on coral reefs form endosymbioses with photosynthetic dinoflagellates in the family Symbiodinaceae, which allows these animals to thrive in oligotrophic habitats (e.g. Baker, 2003; Gates & Edmunds, 1999; Muscatine, McCloskey, & Marian, 1981; Rowan & Powers, 1991; Santos, 2016). In field-collected samples, contamination from symbiodiniaceans is unavoidable, and resulting DNA extractions harbor a mix of anthozoan and dinoflagellete DNA (termed “holobiont DNA”). The combination of slowly evolving mtDNA and dinoflagellate contamination complicates the development of molecular markers suitable for population level questions (e.g. Shearer, Gutiérrez-Rodríguez, & Coffroth, 2005). No broadly useful phylogeographic markers have ever been developed for anthozoans, and thus, most population genetic studies of tropical anthozoans rely on species-specific microsatellite loci to make population-level inferences (e.g. Andras, Rypien, & Harvell, 2013; Baums, Miller, & Hellberg, 2005; Foster et al., 2012; Rippe et al., 2017; Titus et al., 2017a).

The generation of datasets targeting thousands of single nucleotide polymorphisms (SNPs) from anonymous loci via high-throughput sequencing is affordable and provides genome-scale data for non-model organisms. However, marine scientists interested in studying symbiotic anthozoans must still contend with symbiodiniacean contamination in genomic sequence data because there are no simple or reliable ways to completely separate symbiont and host DNA before sequencing. For studies using transcriptomic approaches, anthozoan and dinoflagellate DNA can be parsed bioinformatically, as assembled contigs are long, and conserved, enough to map to published genomic resources (e.g. Davies, Marchetti, Ries, & Castillo, 2016; Kenkel & Matz, 2016; Kenkel, Moya, Strahl, Humphrey, & Bay, 2018). However, the reduced representation sequencing approaches most commonly used in population-level phylogeographic studies (e.g. RADseq, GBS) produce anonymous loci, have short read lengths (e.g. 50-100 bp), and are expected to be recovered largely from non-coding regions. Thus, currently available anthozoan reference genomes will be of limited use to separate dinoflagellate from anthozoan SNPs bioinformatically unless the reference species is closely related to the focal taxa. Likewise, the currently available genomic resources for Symbiodinaceae are also of limited use for parsing reduced representation SNP datasets because of the genetic diversity within its members: long considered to belong to a single genus (*Symbiodinium*), the photosymbiotic dinoflagellates are now recognized to represent 7-15 genus-level lineages (LaJeunesse et al., 2018), with genetic distances between many of these on par with order-level divergences (LaJeunesse et al., 2018; Rowan & Powers, 1992; Santos, 2016). Thus, any symbiodiniacean reference genome used in an attempt to disambiguate endosymbiont and host DNA needs to be very closely related to the specific endosymbiotic dinoflagellate found within the focal anthozoan species to effectively identify dinoflagellate sequences within reduced representation datasets.

Because of the complexity of disentangling host and symbiont sequences from reduced representation sequencing of holobiont DNA, these sequencing approaches have been applied in only limited ways to a small number of photosymbiotic anthozoan species. The majority of these studies come from the scleractinian coral genus *Acropora* (e.g. Devlin-Durante & Baums, 2017; Drury et al., 2017; Shinzato, Mungpakdee, Arakaki, & Satoh, 2015; Rosser et al., 2017;), which have circumvented symbiont contamination by mapping RADseq or GBS loci to the congeneric *Acropora digitifera* reference genome (Shinzato et al., 2011). Others have mapped anonymous loci to conspecific or congeneric transcriptomes and used only the resulting protein-coding SNP datasets for interspecific phylogenetic reconstruction and hybridization studies (e.g. Combosch & Vollmer 2015; Forsman et al., 2017; Johnston et al., 2017). One study employed a subtraction library approach, spinning down homogenized tissue in an effort to remove dinoflagellate cells prior to DNA extraction and creating a separate reduced representation dinoflagellate reference library (Bongaerts et al., 2017). Leydet, Grupstra, Coma, Ribes, & Hellberg (2018) targeted anthozoan RADseq loci by including a congeneric, aposymbiotic, species in their library prep and sequencing- acting as a *de facto* reference library. Each of these studies recognized the importance of removing symbiodiniacean sequences from their reduced representation datasets, acknowledging that successful interpretation of patterns or population parameters requires knowing the extent to which each organism is contributing to the observed patterns.

While true in theory, in practice, how important is it to account for and remove 100% of endosymbiotic dinoflagellate loci from reduced representation datasets? Are reference genomes, or other approaches, always required in order to obtain anthozoan datasets that lead to robust phylogeographic inference and that do not lead spurious results? We hypothesize that, in many commonly encountered circumstances, the unique combination of anthozoan biology, diversity of the endosymbionts, and the manner in which *de novo* SNP-calling programs (i.e. pyRAD, *Stacks*) identify orthologous loci in reduced representation datasets will alleviate the need for anthozoan reference genomes to separate anthozoan from dinoflagellate DNA. This hypothesis rests on several observations. First, many tropical anthozoans have flexible associations that involve diverse lineages of Symbiodiniaceae (e.g. Santos, 2016; Silverstein, Correa, & Baker, 2012). Members of the same host species can harbor different lineages of Symbiodiniaceae (previously called Clades and Types of *Symbiodinium*) in different habitats and across broad geographic space, and even within the same individual or colony (Baker, 2003; Silverstein et al., 2012; Santos, 2016). Second, as outlined above, the genetic divergences between members of Symbiodiniaceae are comparable to order-level differences seen in other dinoflagellates, representing divergences as old as the mid-Jurassic (LaJeunesse et al., 2018, Santos, 2016). Thus, given an adequate sampling distribution, many reduced representation datasets produced for tropical anthozoans will harbor multiple lineages of Symbiodiniaceae. Finally, reduced representation datasets should be comprised of largely short, non-coding DNA fragments. When *de novo* SNP-calling programs cluster DNA sequence fragments and call orthologous loci, the user specifies a missing data threshold before a locus is incorporated into a final dataset. A moderately conservative missing data threshold may be enough to filter out the majority of symbiodiniacean sequences because the program cannot find enough mutationally-conserved, orthologous loci present across the genetically divergent Symbiodiniaceae to meet the missing data thresholds. For example, imagine a RADseq dataset consisting of 10 individuals of an anthozoan from Florida that harbor *Symbiodinium* (previously “Clade A”), and 10 individuals from Bermuda that harbor *Breviolum* (previously *Symbiodinium* “Clade B”), The genetic divergence between *Symbiodinium* and *Breviolum* is large enough (pairwise distance for LSU DNA = 0.37, estimated divergence ~170 mya: LaJeunesse et al., 2018) that few (if any) non-coding orthologous DNA sequences from *Symbiodinium* and *Breviolum* would be retained under a pyRAD missing data threshold requiring a locus to be present in 75% of all individuals. Thus, the loci that would be retained in the final dataset would primarily be from the host anthozoan, which represents intraspecific diversity at shallower evolutionary timescales.

To test this, we used double digest restriction-site associated DNA sequencing (ddRADseq) to reconstruct the range-wide phylogeographic history of the corkscrew sea anemone *Bartholomea annulata*- a species known to harbor multiple members of Symbiodiniaceae throughout the Tropical Western Atlantic (TWA) (see Grajales, Rodríguez, Thornhill, 2016). We then leverage the genome of the sea anemone *Exaiptasia diaphana* (previously *Aiptasia pallida*; see Grajales & Rodríguez, 2014; ICZN, 2017) (Baumgarten et al., 2015), a closely related species from the same family (Aiptasiidae; Grajales & Rodriguez, 2016), to create an aposymbiotic SNP dataset for *B. annulata*. We compare the spatial genetic structure of the aposymbiotic and full holobiont SNP datasets throughout the region, and use coalescent simulation and model selection to understand whether the two datasets are similarly structured and whether they are interpreted as having similar patterns of demographic history with the same parameter estimates (i.e. effective population size, migration) as the aposymbiotic data. We discuss the competing phylogeographic reconstructions and their implications for future studies on symbiotic anthozoans, and further our understanding of the phylogeographic history of Caribbean coral reef taxa.

## 2. Methods

### 2.1. Focal taxon

The corkscrew sea anemone, *B. annulata*, is the most abundant large-bodied species of anemone on coral reef habitats throughout the TWA (Briones-Fourzán, Pérez-Ortiz, Negrete-Soto, Barradas-Ortiz, & Lozano-Álvarez, 2012). Like many tropical anthozoans, it is symbiotic with multiple members of Symbiodiniaceae throughout its range- *Symbiodinium* (formerly Clade A) in Bermuda and Florida and *Cladocopium* (formerly Clade C) in Florida, Mexico, and Panama (Grajales et al., 2016). Symbiodiniaceaens are obtained horizontally after planktonic larvae metamorphose and settle to the benthos, or vertically during pedal laceration. Sexual reproduction occurs twice per year, but asexual reproduction occurs year round (Jennison, 1981).

Although the contribution of sexual and asexual reproduction can vary by habitat type, *B. annulata* appears to rely primarily on sexual reproduction (Titus et al., 2017a). This species also appears to have highly dynamic populations with rapid turnover and a maximum estimated lifespan of ~2 years (O’Reilly & Chadwick, 2017; O’Reilly, Titus, Nelsen, Ratchford, & Chadwick, In Press).

Ecologically, *B. annulata* serves as an important host to the most diverse community of crustacean ectosymbionts of any TWA sea anemone (Briones-Fourzán et al., 2012; Titus & Daly, 2017), including cleaner shrimps that remove parasites from more than 20 families of reef fishes (Huebner & Chadwick, 2012a, b; Titus, Daly, & Exton, 2015a, b; Titus, Vondriska, & Daly, 2017b, Titus, Palombit, & Daly, 2017c). Thus, this species forms the hub of a complex multilevel symbiosis that has potentially radiating effects across multiple trophic levels on TWA reef systems. Finally, this species and its crustacean symbionts are collected commercially by the ornamental aquarium trade along the Florida Reef Tract and are listed by the Florida Fish and Wildlife Conservation Commission as “biologically vulnerable” and “species of conservation concern.”

### 2.2. Sample collection, DNA isolation, library preparation, and data processing

Tissue samples (i.e. tentacle clippings and whole animals) were collected using SCUBA from 14 localities encompassing the entire geographic range of *B. annulata*, and from localities separated by known phylogeographic barriers (Table 1; Fig. 1; Table S1; reviewed by DeBiasse, Richards, Shivji, & Hellberg, 2016). Samples were collected by hand from coral reef habitats between 5- and 15-m depth and preserved on shore using RNAlater. 20-30 samples were collected per locality and transferred back to The Ohio State University for DNA extraction, library preparation, and sequencing. Genomic DNA was isolated using DNeasy Blood and Tissue Kits (Qiagen Inc.) and stored at −20°C. DNA degradation was assessed for each sample using gel electrophoresis, and only samples with high molecular weight DNA were carried forward for ddRADseq library preparation. DNA concentrations were quantified (ng/uL) using a Qubit 2.0 (ThermoFisher) fluorometer and dsDNA broad-range assay kits. 20uL aliquots, each with 200ng of DNA, were prepared for each sample and used for ddRADseq library preparation.

**Table 1.**
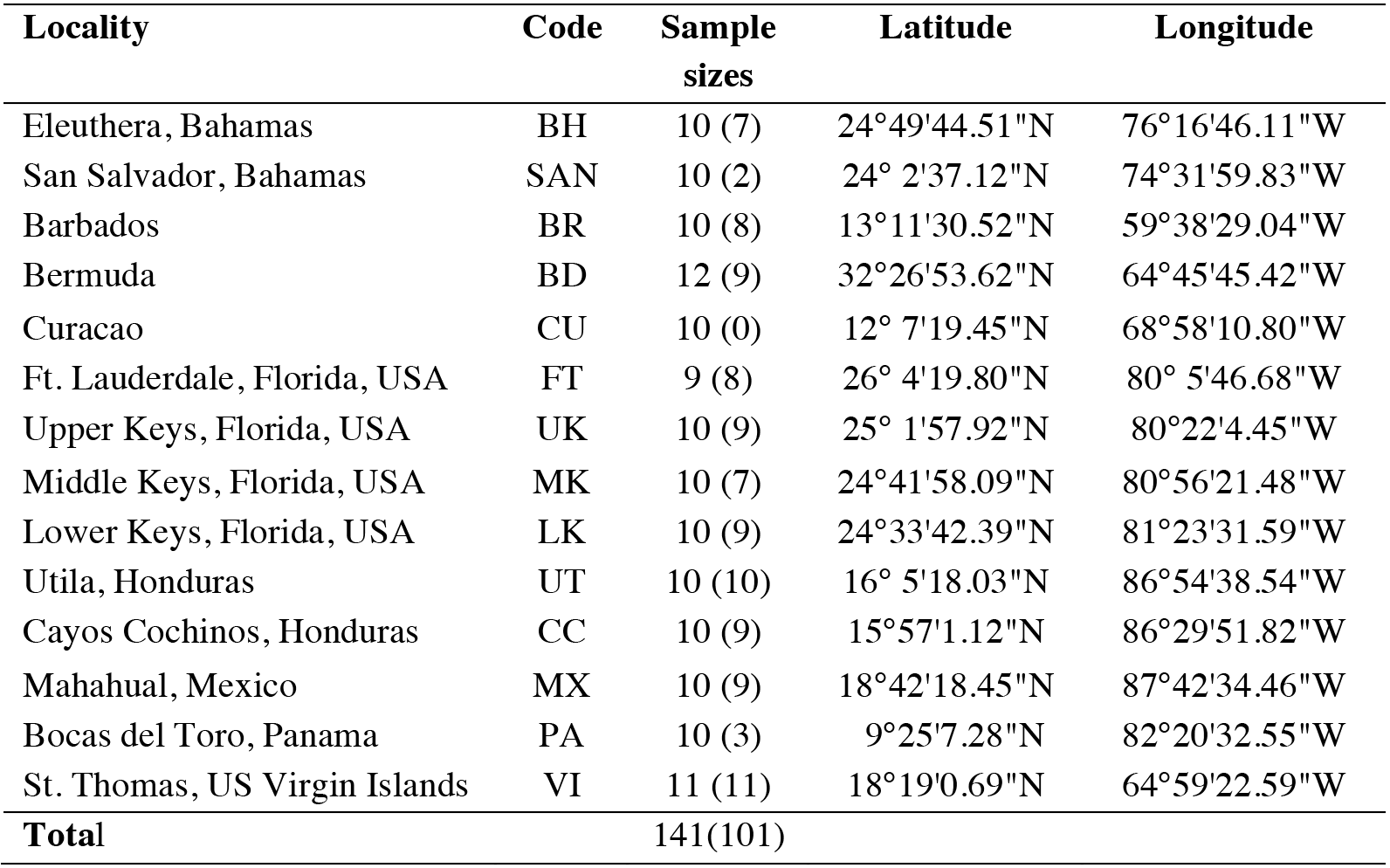
Sample localities, sample sizes, and geographic coordinates of corkscrew sea anemone *Bartholomea annulata* used in this study. Sample sizes reflect the number of samples sequenced and the number of samples retained in the final double digest Restriction-site Associated DNA sequencing (ddRADseq) dataset after accounting for low sequence reads and cryptic species diversity (in parentheses). Differences between the number of samples sequenced and retained reflects variation in the number of sequence reads and sequencing coverage in our ddRADseq dataset across all individuals.

| Locality | Code | Sample sizes | Latitude | Longitude |
| --- | --- | --- | --- | --- |
| Eleuthera, Bahamas | BH | 10 (7) | 24°49'44.51"N | 76°16'46.11"W |
| San Salvador, Bahamas | SAN | 10 (2) | 24° 2'37.12"N | 74°31'59.83"W |
| Barbados | BR | 10 (8) | 13°11'30.52"N | 59°38'29.04"W |
| Bermuda | BD | 12 (9) | 32°26'53.62"N | 64°45'45.42"W |
| Curacao | CU | 10 (0) | 12° 7'19.45"N | 68°58'10.80"W |
| Ft. Lauderdale, Florida, USA | FT | 9 (8) | 26° 4'19.80"N | 80° 5'46.68"W |
| Upper Keys, Florida, USA | UK | 10 (9) | 25° 1'57.92"N | 80°22'4.45"W |
| Middle Keys, Florida, USA | MK | 10 (7) | 24°41'58.09"N | 80°56'21.48"W |
| Lower Keys, Florida, USA | LK | 10 (9) | 24°33'42.39"N | 81°23'31.59"W |
| Utila, Honduras | UT | 10 (10) | 16° 5'18.03"N | 86°54'38.54"W |
| Cayos Cochinos, Honduras | CC | 10 (9) | 15°57'1.12"N | 86°29'51.82"W |
| Mahahual, Mexico | MX | 10 (9) | 18°42'18.45"N | 87°42'34.46"W |
| Bocas del Toro, Panama | PA | 10 (3) | 9°25'7.28"N | 82°20'32.55"W |
| St. Thomas, US Virgin Islands | VI | 11 (11) | 18°19'0.69"N | 64°59'22.59"W |
| **Total** |  | 141(101) |  |  |

**Figure 1.**
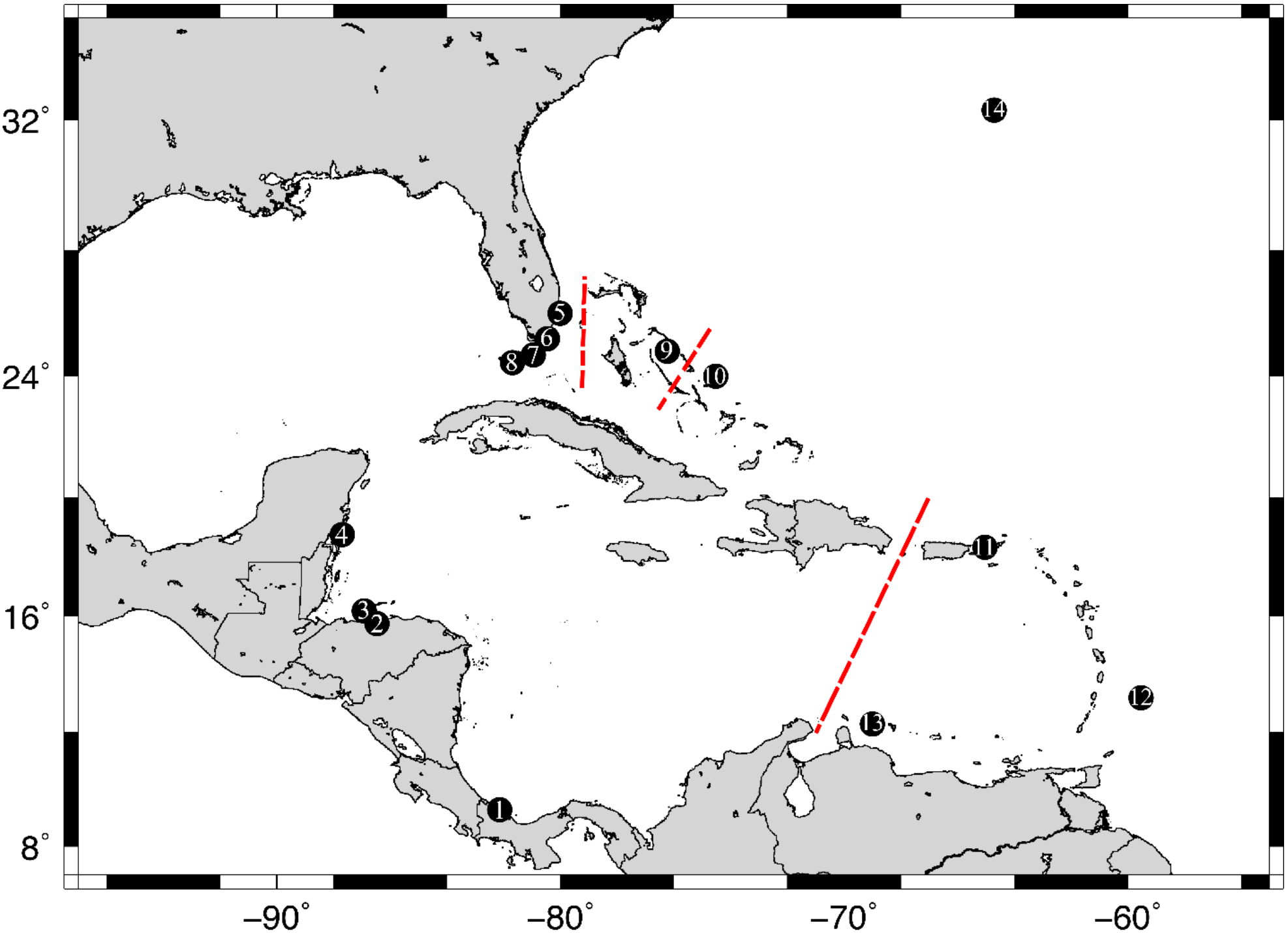
Map of sampling localities throughout the Tropical Western Atlantic for the populations of corkscrew sea anemone *Bartholomea annulata* studied here. 1. Bocas del Toro, Panama, 2) Cayos Cochinos, Honduras, 3) Utila, Honduras, 4) Mahahual, Mexico, 5) Ft. Lauderdale, Florida, 6) Upper Keys, Florida, 7) Middle Keys, Florida, 8) Lower Keys, Florida, 9) Eleuthera, Bahamas, 10) San Salvador, Bahamas, 11) St. Thomas, US Virgin Islands, 12) Barbados, 13) Curacao, 14) Bermuda. Red dashed lines denote previously recovered major phylogeographic breaks in the region.

Between 12-15 individual *B. annulata* samples per locality were carried forward for ddRADseq library preparation. Genomic DNA was digested using two restriction enzymes (*Eco*RI-HF and *psti*-HF), Illumina compatible barcodes were annealed to restriction cut sites, samples were size selected manually using a 400-800 bp size range, and then cleaned using Nucleospin Gel and PCR clean up kits (Macherey-Nagel). Following size selection, each individual sample was amplified using polymerase chain reaction (PCR), cleaned using AMpure XP beads (Agilent), and then quantified via quantitative PCRs (qPCR) to inform the pooling of individual samples into final libraries. A total of 141 individuals (Table 1) met all quality control steps and were pooled across five separate libraries. Samples were sequenced on an Illumina HiSeq 2500 using single-end 100 base pair reads at The Ohio State University Genomics Shared Resource.

### 2.3. Data processing and aposymbiotic dataset assembly

Raw sequence reads were demultiplexed, aligned, and assembled *de novo* using the program pyRAD v3.0.66 (Eaton, 2014). We required a minimum base call Phred score of 20 and set the maximum number of bases in a locus with Phred scores < 20 (NQual) to five. Low quality base calls were replaced with Ns. We set the clustering threshold (Wclust) to 0.90 to assemble reads into loci, and required a minimum coverage depth of seven to call a locus (Mindepth). Finally, we required a locus to be present in 75% of all individuals to be retained in the final dataset. RADseq protocols are known to be susceptible to missing data due to mutations in restriction cut sites and allelic dropout (e.g. Arnold, Corbett-Detig, Hartl, & Bomblies, 2013), but biases can also arise when datasets are overly conservative (i.e. no missing data allowed; Huang & Knowles, 2014). Thus we allowed some missing data in our final dataset.

Previously, we delimited two cryptic lineages of *B. annulata* co-distributed throughout the range in our ddRADseq dataset (Titus, Blischak, & Daly, 2018). After running pyRAD to completion, members of the infrequently sampled lineage (Clade 1, N = 18 individuals; Titus et al., 2018) and individuals with low sequencing coverage (< 500,000 reads; N = 21 individuals), were removed from the dataset, leaving only the more well sampled lineage (Clade 2; n = 101 individuals; Table 1; Table S1). After accounting for cryptic diversity this initial SNP dataset represented our “holobiont dataset,” sequences that were, putatively, a combination of anemone and algal DNA.

To create an anemone-only “aposymbiotic dataset,” we mapped polymorphic loci from the holobiont ddRADseq dataset to the genome of the closely related *Exaiptasia diaphana* (Baumgarten et al., 2015) to identify anemone-only sequences. *Exaiptasia diaphana* and *B. annulata* are members of the same family and are closely related (Grajales & Rodriguez, 2016), and polymorphic microsatellites have previously been designed from *E. diaphana* that amplify in *B. annulata* (Titus et al., 2017a). To map polymorphic *B. annulata* loci to *E. diaphana*, we downloaded the *E. diaphana* genome and created a local BLAST database. After initially running pyRAD to completion, a python script (parse_loci.py, available on Dryad doi:XXX) was written to select the first DNA sequence from each locus in the.loci output file, and create a.fasta file that could then be BLAST-ed against the *E. diaphana* genome (BLAST+; Camacho et al., 2009). We used an 85% identity threshold to call a locus as putatively anemone in origin. Next, a separate python script (blast2loci.py, available on Dryad doi:XXX) was used to read through the BLAST output file, pull all sequences in all loci that met the 85% identity threshold, and create a new.loci file with the same file name as the original. The original.loci file was then replaced with the new anemone-only file, at which point the final step of pyRAD (step 7) was rerun to create our final anemone-only output files (i.e. unlinked SNPs and alleles files) for downstream analyses. As a final check, we created additional local BLAST databases by downloading publicly available endosymbiotic dinoflagellate genomes: *Symbiodinium micradriaticum* (Aranda et al., 2016 as *Symbiodinium micradriaticum* “Clade A”), *Breviolum minutum* (Shoguchi et al., 2013 as *Symbiodinium minutum* “Clade B”), *Cladocopium goreaui* (Liu et al., 2018, as *Symbiodinium goreaui* “Clade C”), and *Fugacium kawagutii* (Liu et al., 2018, as *Symbiodinium kawagutii* “Clade F”). We mapped both our holobiont and apoysymbitic datasets to the symbiodiniacean genomes to see if we could 1) identify any symbiodiniacean sequences in the holobiont data, and 2) confirm that no loci in our aposymbiotic dataset mapped to both symbiodiniacean and *Exaiptasia* genomes. Lastly, we mapped our holobiont dataset to the genome of the distantly related starlet sea anemone, *Nematostella vectensis* (Putnam et al., 2007), to gauge the extent to which intra-order (Actiniaria) genomic resources could be used to effectively identify anemone-only 100bp ddRADseq loci. All scripts for mapping and parsing anemone from symbiodiniacean DNA, along with full details and instructions for using them, can be found on Dryad (doi:XXX).

### 2.4. Population genetic structure

We used the clustering program Structure v2.3.4 (Pritchard, 2000) to infer population genetic structure across the Tropical Western Atlantic. For both holobiont and aposymbiotic datasets, we collapsed bi-allelic data into haplotypes at each locus, thus using information contained in linked SNPs when more than one SNP was present in a locus. Structure analyses were conducted using the admixture model and correlated allele frequencies. Each MCMC chain for each value of *K* was run with a burnin of 1 × 10^5^ generations and sampling period of 2 × 10^5^ generations.

We initially conducted two separate Structure analyses for both the holobiont and aposymbiotic datasets. First, we conducted three iterations of a broad range of *K* values (1- 6) to gain an initial snapshot of the data across the region. In both initial analyses we used the peak ln Pr(D|K) and the Δ*K* (Evanno et al. 2005) to inform the selection of the best *K* value. We then reran Structure using a narrower range of *K* values (1-4) but with more iterations (n = 10). Each MCMC chain for each value of *K* was run with a burnin of 1 × 10^5^ generations and sampling period of 2 × 10^5^ generations. Again, we used ln Pr(D|K) and Δ*K* to select the best value of *K*.

We conducted an analysis of molecular variance (AMOVA) in Arlequin v.3.5 (Excoffier & Lischer, 2010) to test for hierarchical partitioning of genetic diversity across the region. Following our Structure results (see Results), we partitioned samples into Eastern and Western regions. We tested for hierarchical structure among sample localities (*φ*_ST_), among sample localities within a region (*φ*_SC_), and between regions (*φ*_CT_). Calculations in Arlequin v3.5 were made using haplotype data and distance matrices calculated using the number of different alleles per locus. Statistical significance was assessed with 10,000 permutations. Pairwise *φ*_ST_ values were calculated to test for differentiation among sample localities. Genetic diversity summary statistics and pairwise *φ*_ST_ values were also calculated in Arlequin for all sample localities. All calculations were conducted for both aposymbiotic and holobiont datasets.

### 2.5. Demographic modeling selection and parameter estimation

While broad-scale patterns of spatial genetic structure may be robust to some levels of dinoflagellate contamination in reduced representation sequencing datasets, we expect that demographic model selection approaches that make inferences regarding patterns of demographic history, and that generate important population parameter estimates (i.e. effective population sizes, migration rates), should be highly sensitive to the incorporation of data from taxa with different evolutionary histories. Thus, we conducted model selection using the allele frequency spectrum (AFS) and coalescent simulations in the program *fastsimcoal2* (FSC2; Excoffier et al., 2013). FSC2 uses coalescent simulations to calculate the composite likelihood of arbitrarily complex demographic models under a given AFS. The best fit model can then be selected using Akaike information criterion (AIC). We developed 12 user-specified demographic models (Figure 2), all variants of a two-population isolation-migration model as Structure delimited *K* = 2 as the best clustering scheme (see Results). Models differed in the directionality of gene flow, population size changes following divergence, and whether they exhibit patterns of secondary contact following divergence. Genetic clusters in Structure were largely partitioned East and West in the TWA, and 25 individuals from each putative population (50 individuals total; Table S2) were selected to generate two-population, joint-folded, AFS. We conducted model selection on both aposymbiotic and holobiont datasets.

**Figure 2.**
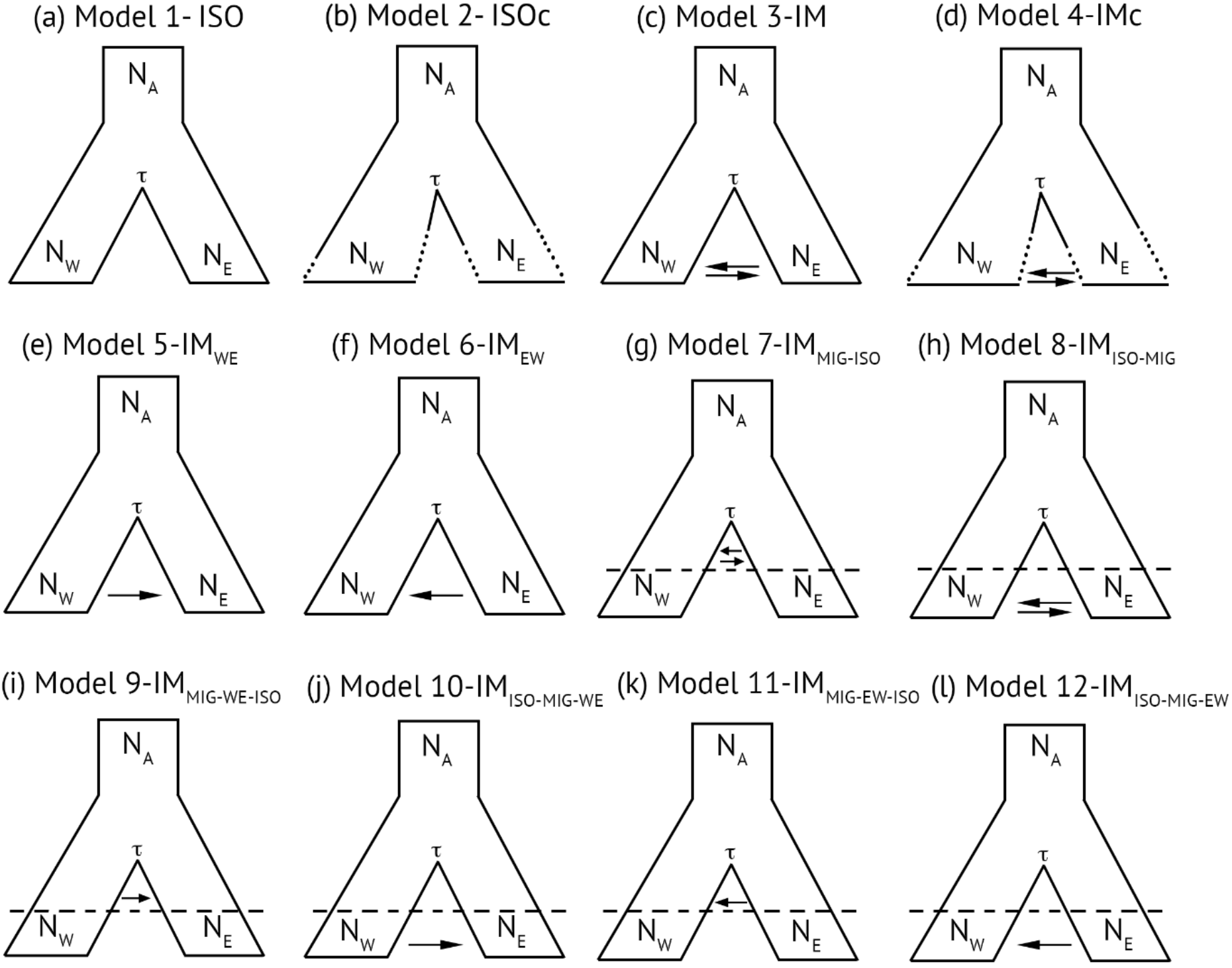
Models used in FSC2 to understand the demographic processes leading to the two-population pattern of diversification in the corkscrew anemone *Bartholomea annulata* across the Tropical Western Atlantic. Each model is a two-population isolation-migration (IM) model that varies in the degree and directionality of gene flow and effective population size. Models are as follows: a) isolation only, b) isolation with population size changes following divergence, c) IM model with symmetric migration, d) IM model with symmetric migration and population size changes, e) IM model with migration from the Western to Eastern population, f) IM model with migration from population Eastern to Western, g) IM model with symmetric migration between populations immediately following divergence followed by more contemporary isolation, h) IM model with isolation immediately following divergence followed by secondary contact and symmetric migration, i) IM model with migration from population Western to Easter immediately following divergence followed by more contemporary isolation, j) IM model with isolation immediately following divergence followed by secondary contact and migration from population Western to Eastern, k) IM model with migration from population Eastern to Western immediately following divergence followed by more contemporary isolation, and l) IM model with isolation immediately following divergence followed by secondary contact and migration from population Eastern to Western.

Two-population, joint-folded AFS were generated from pyRAD output files and previously published python scripts (see Satler & Carstens, 2017). One of the assumptions of FSC2 is that SNPs are in linkage equilibrium (Excoffier et al., 2013), and thus, only one SNP per locus was selected to produce the AFS. Further, AFS calculations in FSC2 require fixed numbers of alleles from all populations (i.e. no missing data). As meeting this latter requirement would greatly decrease our dataset size, and thus likely bias our analyses, we followed the protocol of Satler and Carstens (2017) and Smith et al., (2017) by requiring a locus in our AFS to be present in 85% of all individuals. To account for missing data without violating the requirements of the AFS we built our AFS as follows: 1) if a locus had fewer alleles than our threshold it was discarded, 2) if a locus had the exact number of alleles as the threshold, the minor allele frequency was recorded, and 3) if a locus exceeded the threshold, alleles were down-sampled with replacement until the number of alleles met the threshold, at which point the minor allele frequency was counted. This approach allowed us to maximize the number of SNPs used to build the AFS, but also has the potential to lead to monomorphic alleles based on the down-sampling procedure (see Satler & Carstens 2017). Thus, we repeated the AFS building procedure 10 times, allowing us to account for variation in the down-sampling process during model selection, and also allowing us to calculate confidence intervals on our parameter estimates (Satler & Carstens, 2017; Smith et al., 2017).

Each simulation analysis in FSC2 (i.e. each AFS replicate per model; 12 models × 10 replicates) was repeated 50 times, and we selected the run with the highest composite likelihood for each AFS replicate and model. The best-fit model was then calculated using the AIC and model probabilities calculated following Burnham and Anderson (2002). Because FSC2 requires a per generation mutation rate to scale parameter estimates into real values, we used the substitution per site per generation mutation rate of 4.38 × 10^−8^ proposed for tropical anthozoans (Prada et al., 2017) and a generation time of 1 year for *B. annulata* (Jennison, 1981). All analyses were conducted on the Oakley cluster at the Ohio Supercomputer Center (http://osc.edu).

## 3. RESULTS

### 3.1. Dataset assembly

Double digest RADseq library preparation and sequencing resulted in a total of 186.7 million sequence reads across 141 individuals, 175.6 million of which passed quality control filtering and were retained to create the final dataset. Accounting for individuals with low sequence reads (< 500,000 reads) and cryptic species level diversity (Titus et al., 2018) resulted in a final intraspecific dataset of 101 individuals (Table 1; Table S1). Requiring a locus to be present in a minimum of 75% of all individuals resulted in a final holobiont data set of 11,331 parsimony-informative sites distributed across 3854 unlinked loci. After BLASTing these loci to the *Exaiptasia diaphana* genome, we retained 1402 loci that had matched with high confidence (≥ 85% identity) and these were used as the final aposymbiotic SNP dataset. A total of 59 of the 3854 holobiont ddRADseq loci (~1.5%) mapped to Symbiodiniaceae genomes (Table S3), confirming the presence of at least some symbiont DNA in our holobiont dataset. Of these, 58 mapped to the *S. microadriaticum* genome and one mapped to the *C. goreaui* genome (Table S3). Only five *B. annulata* loci mapped to the starlet sea anemone *N. vectensis* genome (Table S4). SNP files and raw data for both holobiont and aposymbiotic datasets are available on Dryad (Dryad doi:XXX).

### 3.2. Population genetic structure

Genetic clustering analysis in Structure resolved similar patterns across the TWA for both aposymbiotic and holobiont datasets. For the aposymbiotic dataset, *K* = 2 was selected by Structure using both lnP(*K*) and Δ*K* as the best clustering scheme (Fig. 3; Table S5). Diversity was largely binned into Western and Eastern partitions, but with admixture (Figure 3). The most notable genetic break was that between the Lower Keys (LK) and Eleuthera, Bahamas (BH), sample localities in close proximity and bisected by the Florida Straits (Figs. 1 & 3). The holobiont dataset recovered similar geographic partitioning, but Structure selected *K* = 3 as the best partitioning scheme using lnP(*K*) and Δ*K* (Fig. 3; Table S6). The additional genetic cluster did not illuminate any unrecovered geographic partitioning across the region beyond what was recovered by a *K* = 2 partitioning scheme (Fig. 3). The West-East genetic break across the TWA, with admixture, is still largely resolved in the holobiont dataset with the most notable break again between the LK and BH sample localities (Fig. 3b).

**Figure 3.**
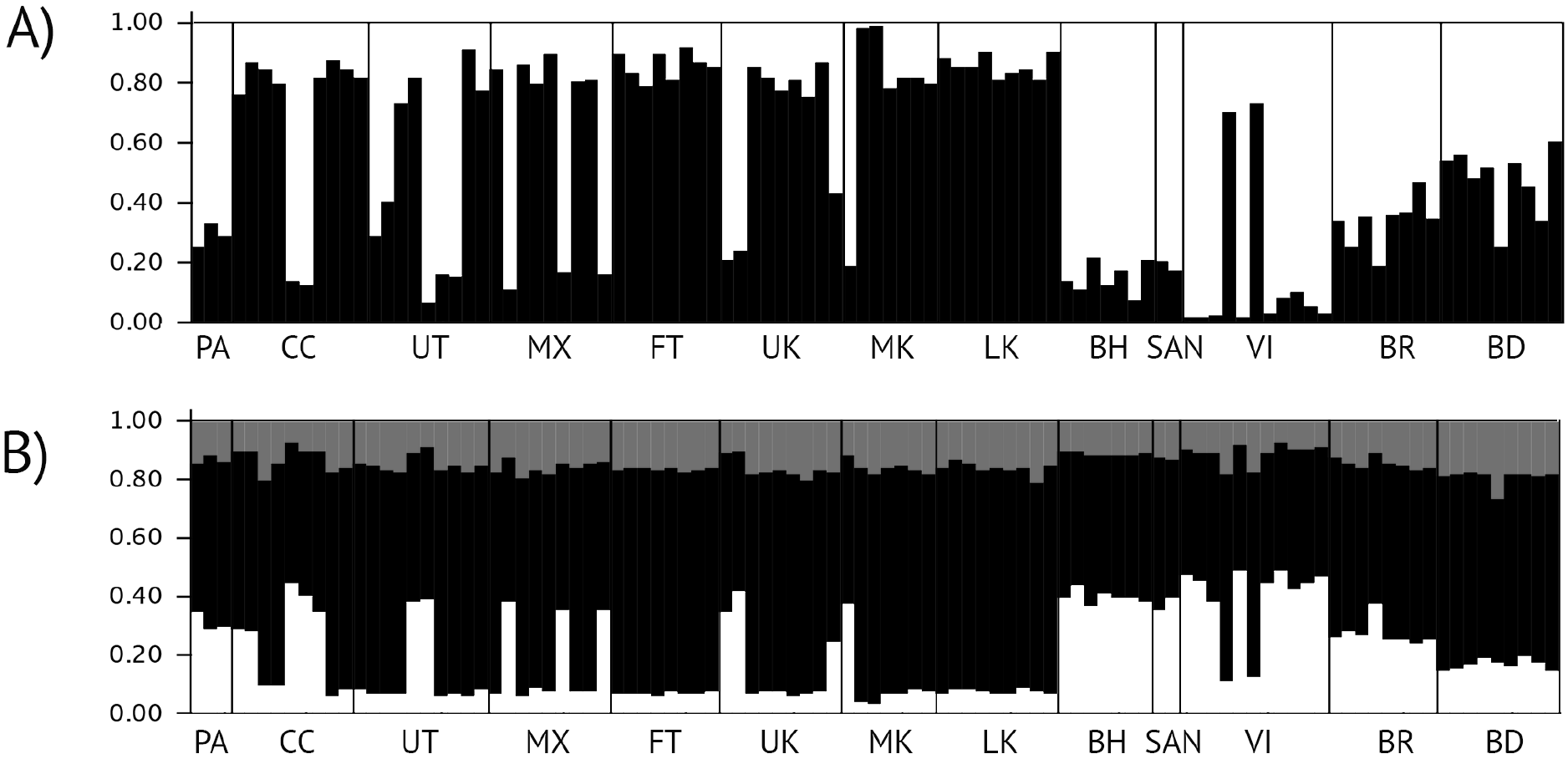
A) Genetic clustering results (*K* =2) for the aposymbiotic *Bartholomea annulata* RADseq dataset. B) Genetic clustering results (*K* = 3) for the holobiont *Bartholomea annulata* RADseq dataset. Samples are partitioned by sample locality in a largely West to East (left to right) geographic layout. 1. Bocas del Toro, Panama, 2) Cayos Cochinos, Honduras, 3) Utila, Honduras, 4) Mahahual, Mexico, 5) Ft. Lauderdale, Florida, 6) Upper Keys, Florida, 7) Middle Keys, Florida, 8) Lower Keys, Florida, 9) Eleuthera, Bahamas, 10) San Salvador, Bahamas, 11) St. Thomas, US Virgin Islands, 12) Barbados, 13) Bermuda.

Population genetic analyses in Arlequin reflect nearly identical results for both datasets. AMOVA results indicate low, but significant, population genetic structure at all hierarchical levels for both datasets, and both datasets have similar patterns of genetic variation at each hierarchical level (Table 2). Similarly, pairwise *φ*_ST_ values calculated by Arlequin were low, but significant, among many sample localities for both datasets (Table 3), and there were no major differences in the genetic diversity summary statistics for both datasets across the range (Table 4).

**Table 2.** Analysis of Molecular Variance (AMOVA) results for aposymbiotic and holobiont *Bartholomea annulata* RADseq datasets. Data were partitioned into Eastern and Western Regions and reflect nearly identical levels of genetic diversity partitioned at all hierarchical levels. **p < 0.0005; *p < 0.005.

| Dataset | Among localities | $\varphi_{ST}$ | Among localities within regions | $\varphi_{SC}$ | Among regions | $\varphi_{CT}$ |
| --- | --- | --- | --- | --- | --- | --- |
| Aposymbiotic | 96.55 | 0.03** | 2.24 | 0.02** | 1.21 | 0.01* |
| Holobiont | 95.90 | 0.04** | 2.62 | 0.03** | 1.48 | 0.01* |

**Table 3.** Pairwise *φ*_ST_ calculated among sample localities for aposymbiotic (below diagonal) and holobiont (above diagonal) *Bartholomea annulata* datasets. Locality codes correspond to those in Figure 1. *φ*_ST_ values highlighted and bolded are significant at p < 0.05. ^denotes *φ*_ST_ values significant in the holobiont dataset only

|  | PA | CC | UT | MX | FT | UK | MK | LK | BH | SAN | VI | BR | BD |
| --- | --- | --- | --- | --- | --- | --- | --- | --- | --- | --- | --- | --- | --- |
| PA | - | **0.05** | **0.03** | **0.08** | **0.05** | **0.03** | **0.04** | **0.04** | **0.05** | **0.03** | **0.03** | **0.03**<sup>^</sup> | **0.03** |
| CC | **0.05** | - | **0.05** | 0.04 | **0.04** | **0.04** | **0.03** | **0.07** | **0.07** | **0.03** | **0.03** | **0.03** | **0.05** |
| UT | **0.03** | **0.04** | - | **0.08** | **0.04** | **0.03** | **0.05** | **0.05** | **0.05** | **0.04** | **0.05** | **0.03** | **0.05** |
| MX | **0.06** | 0.02 | **0.07** | - | **0.07** | **0.05**<sup>^</sup> | **0.09** | **0.09** | **0.08** | **0.07** | **0.08** | **0.05**<sup>^</sup> | **0.07** |
| FT | **0.05** | **0.03** | **0.04** | **0.06** | - | **0.03** | **0.06** | **0.06** | **0.06** | **0.04** | **0.03**<sup>^</sup> | **0.04** | **0.05** |
| UK | **0.03** | **0.04** | **0.02** | 0.03 | **0.02** | - | **0.03** | **0.02** | **0.03** | 0.00 | 0.00 | **0.01**<sup>^</sup> | **0.02** |
| MK | **0.03** | **0.02** | **0.03** | **0.07** | **0.06** | **0.02** | - | **0.02**<sup>^</sup> | **0.03** | 0.00 | 0.02 | 0.01 | **0.02** |
| LK | **0.03** | **0.06** | **0.04** | **0.07** | **0.06** | **0.02** | 0.01 | - | **0.03** | 0.00 | 0.02 | **0.02** | **0.02** |
| BH | **0.04** | **0.06** | **0.05** | **0.05** | **0.06** | 0.02 | **0.02** | **0.03** | - | 0.01 | 0.02 | **0.02** | **0.03** |
| SAN | **0.02** | **0.02** | **0.03** | **0.07** | **0.04** | 0.00 | 0.00 | 0.00 | 0.00 | - | **0.06** | 0.00 | 0.00 |
| VI | 0.02 | **0.04** | **0.05** | **0.04** | 0.04 | 0.00 | 0.00 | 0.01 | 0.02 | **0.07** | - | 0.00 | 0.00 |
| BR | **0.02** | **0.03** | **0.02** | 0.04 | **0.03** | 0.01 | 0.01 | **0.02** | **0.02** | 0.00 | 0.00 | - | **0.02**<sup>^</sup> |
| BD | **0.03** | **0.04** | **0.03** | **0.05** | **0.04** | **0.01** | **0.01** | **0.02** | **0.03** | 0.00 | 0.02 | 0.01 | - |

**Table 4.** Diversity indices calculated from aposymbiotic and holobiont (in parentheses) RADseq data for *Bartholomea annulata* across the Tropical Western Atlantic. Diversity indices calculated for each sample locality and for each genetically defined population grouping determined by Structure. Values reflect nearly identical genetic diversity indices between aposymbiotic and holobiont datasets all sample localities. *N*_*G*_ = Number of gene copies, SS = segregating sites, *θ* = theta calculated from segregating sites, *π* = nucleotide diversity, Ho = observed heterozygosity, He = expected heterozygosity.

| Sample locality | Code | $N_G$ | SS | $\theta$ | $\pi$ | Ho | He |
| --- | --- | --- | --- | --- | --- | --- | --- |
| Eleuthera | BH | 14 | 169<br>(169) | 53.1<br>(53.1) | 0.030<br>(0.028) | 0.20<br>(0.18) | 0.21<br>(0.19) |
| San Salvador | SAN | 4 | 93<br>(84) | 50.7<br>(45.8) | 0.033<br>(0.029) | 0.45<br>(0.48) | 0.52<br>(0.51) |
| Barbados | BR | 16 | 166<br>(140) | 50.02<br>(42.1) | 0.031<br>(0.026) | 0.17<br>(0.17) | 0.19<br>(0.18) |
| Bermuda | BD | 18 | 197<br>(177) | 57.7<br>(51.4) | 0.035<br>(0.032) | 0.15<br>(0.15) | 0.17<br>(0.16) |
| Curacao | CU | - | - | - | - | - | - |
| Ft. Lauderdale, Florida, USA | FT | 16 | 206<br>(178) | 62.08<br>(53.6) | 0.031<br>(0.026) | 0.17<br>(0.15) | 0.18<br>(0.16) |
| Upper Keys, Florida, USA | UK | 18 | 221<br>(198) | 64.2<br>(57.5) | 0.03<br>(0.028) | 0.15<br>(0.14) | 0.16<br>(0.15) |
| Middle Keys, Florida, USA | MK | 14 | 177<br>(151) | 55.6<br>(47.4) | 0.029<br>(0.027) | 0.18<br>(0.19) | 0.20<br>(0.20) |
| Lower Keys, Florida, USA | LK | 18 | 198<br>(169) | 57.56<br>(49.3) | 0.033<br>(0.026) | 0.17<br>(0.16) | 0.18<br>(0.16) |
| Cayos Cochinos, Honduras | CC | 18 | 194<br>(169) | 56.4<br>(49.1) | 0.029<br>(0.025) | 0.15<br>(0.14) | 0.16<br>(0.15) |
| Utila, Honduras | UT | 20 | 179<br>(163) | 50.4<br>(45.9) | 0.027<br>(0.042) | 0.13<br>(0.12) | 0.14<br>(0.13) |
| Mexico | MX | 18 | 152<br>(148) | 44.1<br>(43.0) | 0.033<br>(0.032) | 0.14<br>(0.14) | 0.16<br>(0.15) |
| Panama | PA | 6 | 68<br>(59) | 29.8<br>(25.8) | 0.025<br>(0.022) | 0.33<br>(0.35) | 0.39<br>(0.38) |
| US Virgin Islands | VI | 22 | 152<br>(171) | 41.6<br>(46.9) | 0.026<br>(0.027) | 0.14<br>(0.13) | 0.15<br>(0.13) |

### 3.3. Demographic model selection

Coalescent modeling in FSC2 returned identical model selection results between aposymbiotic and holobiont datasets (Table 5). For both, Akaike Information Critereon (AIC) selected model 6, an IM model with unidirectional gene flow from East to West, as the best-fit model (Fig. 2). According to Akaike model weights, model 6 received over 0.70 of the support (Table 5) in both aposymbiotic and holobiont datasets. A secondary contact model (Model 10; Fig. 2), with isolation immediately after divergence followed by secondary contact and unidirectional West-East gene flow, received the next highest amount of support according to AIC, although the Akaike weight differed in how much support was given to each, with the aposymbiotic dataset having a clearer preference for this model over the next best one, compared to the holobiont data set (Table 5).

**Table 5.** Akaike Information Criterion results for model selection from FSC2 for the aposymbiotic and holobiont (in parentheses) *Bartholomea annulata* datasets. Model rank was identical between aposymbiotic and holobiont datasets, with broadly similar model likelihoods and model weights. Model refers to those depicted and described in Figure 2. *k* = number of parameters in the model, AIC = Akaike Information Criterion, Δ_I_ = change in AIC scores, and *w*_*i*_ = Akaike weights. Models are listed according to their AIC rank and the highest ranked model is highlighted.

| Model | $k$ | ln(Likelihood) | AIC | $\Delta_i$ | Model Likelihoods | $w_i$ |
| --- | --- | --- | --- | --- | --- | --- |
| 6 - IM<sub>EW</sub> | 5 | -8846.5<br>(-19863.5) | 17703.1<br>(39737.0) | 0<br>(0) | 1<br>(1) | 0.75<br>(0.73) |
| 10 - IM<sub>ISO-MIG-WE</sub> | 6 | -8847.3<br>(-19863.5) | 17706.7<br>(39739.0) | 3.5<br>(2.0) | 0.17<br>(0.36) | 0.12<br>(0.26) |
| 5 - IM<sub>WE</sub> | 5 | -8848.4<br>(-19870.0) | 17706.9<br>(39750.1) | 3.7<br>(13.1) | 0.15<br>(0.001) | 0.11<br>(0.001) |
| 3 - IM | 6 | -8853.0<br>(-19877.1) | 17718.0<br>(39766.3) | 14.8<br>(29.3) | $6.0e^{-3}$<br>($4.2e^{-7}$) | $4.0e^{-3}$<br>($3.1e^{-7}$) |
| 8 - IM<sub>ISO-MIG</sub> | 7 | -8857.7<br>(-19844.6) | 17729.5<br>(39783.3) | 26.3<br>(46.3) | $1.8e^{-6}$<br>($8.6e^{-11}$) | $1.4e^{-6}$<br>($6.3e^{-11}$) |
| 1 - ISO | 4 | -8868.3<br>(-19922.8) | 17744.6<br>(39853.7) | 41.4<br>(116.7) | $1.0e^{-9}$<br>($4.5e^{-26}$) | $7.7e^{-10}$<br>($3.3e^{-26}$) |
| 12 - IM<sub>ISO-MIG-EW</sub> | 6 | -8885.5<br>(-19963.2) | 17783.0<br>(39938.4) | 79.8<br>(201.3) | $4.6e^{-18}$<br>($1.8e^{-44}$) | $3.4e^{-18}$<br>($1.3e^{-44}$) |
| 2 - ISOc | 11 | -10965.6<br>(-22778.0) | 21953.3<br>(45578.1) | 4250.0<br>(5841.1) | 0<br>(0) | 0<br>(0) |
| 4 - IMc | 12 | -11569.8<br>(-24493.4) | 23163.7<br>(49010.9) | 5460.5<br>(9272.9) | 0<br>(0) | 0<br>(0) |
| 9 - IM<sub>MIG-WE-ISO</sub> | 6 | -13772.4<br>(-30574.7) | 27556.8<br>(61161.5) | 9853.6<br>(21424.4) | 0<br>(0) | 0<br>(0) |
| 7 - IM<sub>MIG-ISO</sub> | 7 | -13772.0<br>(-30576.1) | 27558.1<br>(61166.2) | 9854.9<br>(21429.1) | 0<br>(0) | 0<br>(0) |
| 11 - IM<sub>MIG-EW-ISO</sub> | 6 | -13947.7<br>(-30985.4) | 27907.5<br>(61982.8) | 10204.3<br>(22245.8) | 0<br>(0) | 0<br>(0) |

Parameter values and confidence intervals for effective population size, divergence time, and migration rate estimated from FSC2 simulations were entirely overlapping between aposymbiotic and holobiont datasets (Table 6). For both datasets, FSC2 estimated that Eastern *B. annulata* populations had greater effective population sizes than Western populations, and that per-generation migration rate was low (Table 6). Divergence time estimates varied more than other parameter values but still had overlapping confidence intervals. The aposymbiotic dataset had an estimated a mean divergence time between Eastern and Western populations at ~39,000 ybp, whereas the holobiont dataset had an estimated a mean divergence time between populations at ~58,000 ybp.

**Table 6.**
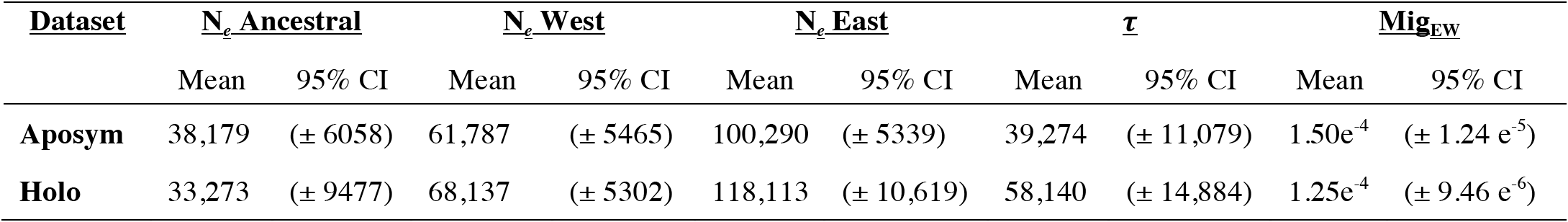
Parameter estimates and 95% confidence intervals (CI) generated from FSC2 coalescent simulations for aposymbiotic (Aposym) and holobiont (Holo) *Bartholomea annulata* datasets. N_*e*_ = effective population size, τ = divergence time, Mig_EW_ = migration rate from Eastern to Western populations. Values reported for N_*e*_ are in number of individuals and the values for τ are reported in years before present. Parameter values reflect overlapping confidence intervals between aposymbtiotic and holobiont for every parameter calculated.

| <u>Dataset</u> | <u>$N_e$ Ancestral</u> |  | <u>$N_e$ West</u> |  | <u>$N_e$ East</u> |  | <u>$\tau$</u> |  | <u>$Mig_{EW}$</u> |  |
| --- | --- | --- | --- | --- | --- | --- | --- | --- | --- | --- |
|  | Mean | 95% CI | Mean | 95% CI | Mean | 95% CI | Mean | 95% CI | Mean | 95% CI |
| **Aposym** | 38,179 | ($\pm 6058$) | 61,787 | ($\pm 5465$) | 100,290 | ($\pm 5339$) | 39,274 | ($\pm 11,079$) | $1.50e^{-4}$ | ($\pm 1.24 e^{-5}$) |
| **Holo** | 33,273 | ($\pm 9477$) | 68,137 | ($\pm 5302$) | 118,113 | ($\pm 10,619$) | 58,140 | ($\pm 14,884$) | $1.25e^{-4}$ | ($\pm 9.46 e^{-6}$) |

## 4. DISCUSSION

### 4.1. Reduced representation sequencing for symbiotic anthozoans

Symbiodiniacean DNA contamination has represented a substantial hurdle for researchers working on tropical anthozoans (e.g. Shearer et al., 2005; Bongaerts et al., 2017; Leydet et al., 2018), and high-throughput sequencing has done little to alleviate these issues in population-level studies. Based on the analyses we conduct here, however, we fail to reject our hypothesis that, anthozoan reference genomes may not always be necessary for making reliable spatial and demographic phylogeographic inference including, importantly, population parameter estimates. Instead, we find broadly similar interpretations from a holobiont dataset and from one “cleaned” of symbiont sequences. Understanding the evolutionary and historical processes that have shaped the diversity of tropical anthozoans has been, and will continue to be, an important research priority for marine phylogeographers (Bowen, Rocha, Toonen, Karl, & ToBo Lab, 2016; Bowen et al., 2016). Our results present a promising framework and way forward for researchers wishing to employ these reduced representation sequencing approaches on symbiotic anthozoan species that are not closely related to species with currently available reference genomes.

The framework and experimental design of our study, effectively a single-species phylogeographic study that spans the entire range of our focal taxon, is representative of many studies that examine the spatial and demographic history of a given species. Although the degree to which symbiotic anthozoans are specific to a particular lineage of Symbiodiniaceae is unresolved, evidence is mounting that these associations are spatially and temporally variable, particularly in stony corals, where much of this research has focused (e.g. Silverstein et al., 2012). Thus, we believe that given a broad sampling scheme with respect to geography and habitat, *de novo* assembly and SNP-calling programs will act as *de facto* filtering programs for symbiodiniaceans in many reduced representation datasets produced from symbiotic anthozoans. Resulting datasets will be overwhelmingly comprised of anthozoan DNA loci.

In our analyses, we would expect that more than doubling of our dataset (~1400 to 3800 SNPs) by the inclusion of putative symbiont loci in our holobiont dataset would lead to major differences in interpretation. The ~2400 uncharacterized loci that did not map to the *E. diaphana* genome represent some combination of anemone and symbiont sequences. Because each anthozoan tentacle cell can contain dozens of *Symbiodinium* cells, and thus *Symbiodinium* nuclei often outnumber anemone nuclei (reviewed by Davy, Allemand, & Weis, 2012), the holobiont dataset could reflect a greater contribution from dinoflagellates than from *B. annulata.* If even only half of the 2400 uncharacterized loci our holobiont dataset were from members of Symbiodiniaceae, we would expect them to greatly influence our holobiont analyses, especially our parameter estimates, which should be the most sensitive to the incorporation of sequence data from multiple species with different evolutionary histories. That we recover indistinguishable phylogeographic histories with completely overlapping diversity indices and parameter estimates leads us to hypothesize that we have very few symbiodiniacean loci in our holobiont dataset. The use of a sea anemone reference genome from the same family rather than a congeneric or conspecific reference genome is the most likely explanation for why ~2300 loci remain uncharacterized in our holobiont dataset: these loci are not shared between *E. diaphana* and *B. annulata*, and so are not included in the aposymbiotic dataset (because that uses the *E. diaphana* genome as a probe for putative anemone loci). Although mapping our reads to genomic resources from members of Symbiodiniaceae confirms we do have some dinoflagellate sequence data in our holobiont dataset (at least ~1.5% of all loci), these are such a small fraction of the SNPs that they may simply be genetic “noise,” swamped out by orders of magnitude more anthozoan SNPs.

Conspecific, or congeneric, reference genomes clearly represent the best approach to removing symbiont loci from reduced representation datasets. However, to date, there are only a handful of published anthozoan genomes (Baumgarten et al. 2015; Prada et al., 2017; Putnam et al. 2007; Shinzato et al. 2011; Wang, Liew, Li, Zoccola, Tambutte, & Aranda, 2017; Voolstra et al., 2017). Our study demonstrates that reference genomes within the same family may serve as adequate genomic resources, but reference genomes that are simply within the same order are likely too distant to serve in the same capacity, at least for actiniarians: only five loci from *B. annulata* (suborder Anthemonae, superfamily Metridioidea, family Aiptasiidae) mapped to the genome of *Nematostella vectensis* (suborder Anenthemonae, superfamily Edwardsioidiea, Family Edwardsiidae). This point parallels the observation that using a reference genome of the endosymbiotic dinoflagellate without concern for the particular lineage of symbiodiniacean harbored by a particular anthozoan is unlikely to remove all dinoflagellate loci.

From a practical standpoint, we recommend that studies employing reduced representation approaches for symbiotic anthozoans without genomic resources from closely related species 1) employ extensive geographic sampling, or sample broadly across ecologically disjunct habitats (i.e. depth, temperature, nutrient concentration) to maximize the likelihood of sampling hosts that harbor diverse symbiodiniaceans and 2) demonstrate empirically that multiple lineages of Symbiodiniaceae are represented in the collected samples via PCR or sequencing (e.g. ITS, cp23s). In host species with highly specific endosymbiont associations, the approach to sampling and sequencing we describe here would likely be ineffective, as orthologous symbiodiniacean loci would be present in all samples and sample localities, and *de novo* clustering programs would not filter these out. In these cases, employing approaches like those of Bongaerts et al. (2017) or Leydet et al. (2018) may be required. Finally, for a host species where population genetic differentiation is driven by a handful of SNPs under selection, incorporating even a small number of symbiont loci could mask important signal. This is unknowable *a priori*, and studies wanting to analyze holobiont DNA at the population level should acknowledge these limitations, and follow up studies should be conducted once reference genomes are available.

### 4.2. *Phylogeographic history of* Bartholomea annulata

The phylogeographic history of coral reef communities in the TWA most often revolves around a major barrier to dispersal at the Mona Passage, separating Hispanola from Puerto Rico (e.g. Baums et al., 2005; Hellberg, 2007; DeBiasse et al. 2016). This barrier has been well resolved for a number of stony corals, fishes, and other invertebrates (reviewed by DeBiasse et al. 2016). Our range-wide phylogeographic analysis demonstrates that the corkscrew sea anemone *Bartholomea annulata* shows subtle, but significant, genetic structure across the TWA, with the Florida Straits, rather than the Mona Passage, being the most well resolved phylogeographic break in the region. Further, while we demonstrate a number of low, but significant, *φ*_ST_ values across many sample localities, genetic clustering loosely groups *B. annulata* into Eastern and Western populations (Fig. 3). The Bahamas and the Florida Keys, sample localities immediately to the East and West of the Florida Straits are separated by ~100 km, but is the region with the clearest genetic partitioning (Figure 3). Sample localities further East (e.g. Barbados, Bermuda) and West (e.g. Mexico, Honduras) exhibit more genetic admixture and may have experienced more historical and contemporary gene flow. No other major genetic structuring was recovered across the TWA, although phylogeographic breaks and regions with unique genetic diversity are known for other groups of organisms, including the Southern Caribbean phylogeographic break between Panama and Curacao, a proposed Central Bahamas phylogeographic break, and regions such as the Meso-American Barrier Reef, Panama, and Bermuda (reviewed by DeBiasse et al. 2016).

Across the Florida Straits, demographic model selection suggests that the best fit for these data among the models we tested is a two-population pattern with continuous unidirectional gene flow from East to West following divergence (Table 5). An important note is that as the coalescent is a backwards-in-time framework, a model with gene flow from East to West reflects forward-in-time gene flow from West to East. This largely fits with the prevailing currents in the TWA, as currents that deflect North in the Western Caribbean basin ultimately form the Loop Current in the Gulf of Mexico and then are forced East through the narrow stretch of sea between Florida and Cuba before turning North again and forming the Gulf Stream. Contemporary gene flow from East to West, would most likely occur in the Southern Caribbean where equatorial currents flow westward near the Southern Windward Islands.

Divergence time estimates from FSC2 between Eastern and Western populations given the current estimate of mutation rate suggests a recent divergence between populations of *B. annulata* sometime within the last 30,000-50,000 years (Table 6), firmly within the most recent glacial maxima (15,000-100,000 years before present). During this time, sea level would have been as much as 120 m below present day levels, and both the Florida peninsula and the Bahamas platform would have been sub-aerially exposed, significantly increasing the amount of dry land subdividing the region and also decreasing available reef habitat (reviewed by Ludt & Rocha 2015). This would have been especially true for Eleuthera, which would have been isolated from the Florida Straits by two large portions of the then-dry-land Bahamas Banks, and two enclosed deep water trenches (Tongue of the Ocean and the Exuma Sound). Water exchange, and thus potential for dispersal and gene flow, would have been greatly reduced during this period, allowing for allopatric divergence and local retention of larvae. This scenario would fit well with a phylogeographic model of divergence followed by a period of isolation, then secondary contact and migration during more recent interglacial periods which coincided with sea level rise. A secondary contact model was the next best fit to our data according to AIC (Table 5). However, we either 1) do not have enough signature in the data for it to be selected as the best fit, or, 2) even though the Florida and Bahamas populations would have been largely isolated, other sample localities in the Eastern (i.e. Virgin Islands, Barbados) and Western (i.e. Mexico, Honduras) would not have been isolated to the same extent, and gene flow between these localities could be responsible for the unidirectional gene flow we see in our best-fit models.

Interestingly, FSC2 simulations and population summary statistics estimate larger effective population sizes in the Eastern Caribbean than in the West (Table 6). At face value, this seems to be at odds with the current geography of the Western Caribbean basin as there is more submerged continental-shelf shallow-water habitat in the Western Caribbean and Florida than there is in the Eastern Caribbean (Ludt & Rocha, 2015), where coral reef habitat is largely limited to small fringing reefs around islands of volcanic origin. Unidirectional gene flow from the West to the East, as recovered by our best-fit model, could be responsible for this increase in effective population size, with the Eastern Caribbean effectively a sink of a genetic diversity. In addition, the Bahamas are a large, shallow, archipelago and likely capable of supporting immense census population sizes of *B. annulata.* As a habitat generalist, *B. annulata* can colonize hard bottom, seagrass, mangrove, and coral-dominated habitats (e.g. Briones-Fourzán et al. 2012; O’Reilly & Chadwick, 2017; Titus et al., 2017a), and is thus, not limited strictly to fore reefs. Large habitat space with genetic input from Western population could be driving this pattern.

### 4.3. Conclusions

Our study demonstrates that the corkscrew sea anemone, *Bartholomea annulata*, exhibits weak genetic structure across the Tropical Western Atlantic, and that demographic modeling of this species suggests that unidirectional gene flow from the western to eastern Caribbean can largely explain the observed patterns of genetic diversity. Interestingly, we recover the same spatial and demographic patterns, including entirely overlapping parameter estimates, regardless of whether we use an aposymbiotic ddRADseq dataset or whether we use our putative holobiont dataset. Although we can confirm that at least ~1.5% of the loci in the holobiont dataset are from members of Symbiodiniaceae, we hypothesize that the remaining ~2400 uncharacterized loci are primarily from *B. annulata*, representing SNPs not shared with *Exaiptasia diaphana*. Because of the diversity of dinoflagellate lineages hosted by *B. annulata* across its range and the genetic divergence within and among lineages within Symbiodiniaceae, we believe the manner in which *de novo* reduced representation clustering algorithms assemble RADseq datasets effectively removes most of the SNPs from the photosymbionts.

To further test our hypothesis, this study should be repeated with an anthozoan species with flexible *Symbiodinium* associations and that has a conspecific reference genome available (e.g.,in *Exaiptasia diaphana*, *Acropora digitifera*, *Stylophora pistillata*). This would allow the exact number of anthozoan SNPs identified in the final dataset to be quantified rather than leaving a large fraction of SNPs uncharacterized. Nonetheless, our findings represent an important avenue along which future research on symbiotic anthozoans can continue until greater numbers of reference genomes can be sequenced, annotated, and made publicly available. Tropical anthozoans form the foundation of ecosystems that rival rainforests in diversity, perform important ecological roles, have commercial value, and are especially vulnerable to climate change (e.g. Hughes et al. 2017; Palumbi et al., 2014). As selection can act rapidly on standing genetic diversity (Przeworski et al., 2005; Barrett & Schluter, 2008; Reid et al., 2016), understanding the historical processes that have shaped contemporary distributions of diversity can help set conservation priorities in a rapidly changing climate.

## Acknowledgements

We are grateful to Erich Bartels, Annelise del Rio, Jose Diaz, Dan Exton, Lisle Gibbs, Natalie Hamilton, Alex Hunter, Anna Klompen Jason Macrander, Spencer Palombit, Stephen Ratchford, Nancy Sheridan Nuno Simoes, Jill Titus, Cory Walter, Eric Witt, Clay Vondriska, and the Operation Wallacea dive staff for assistance in the field and laboratory. We also thank Paul Blischak, Jordan Satler, Megan Smith, and Bryan Carstens for bioinformatic assistance and advice regarding *fastsimcoal2* analyses and model selection. Bellairs Research Station, the Bermuda Institute of Ocean Science, Cape Eleuthera Institute, CARMABI, Coral View Dive Center, Gerace Research Centre, the Honduran Coral Reef Foundation, Mote Marine Laboratory, Smithsonian Tropical Research Institute, and the University of the Virgin Islands provided valuable logistical support in the field. Specimens were collected from throughout the Tropical Western Atlantic under permits: SE/A-88- 15, PPF/DGOPA-127/14, CZ01/9/9, FKNMS-2012-155, SAL-12-1432A-SR, STT037-14, 140408, MAR/FIS/17, and 19985. This research was supported by funding from a National Science Foundation-Doctoral Dissertation Improvement Grant DEB-1601645 and Florida Fish and Wildlife Conservation Commission awards to B.M.T. & M.D. Operation Wallacea, American Philosophical Society, International Society for Reef Studies Graduate Fellowship, PADI Foundation Grant, and American Museum of Natural History Lerner Gray Funds supported field research for B.M.T. Additional funding was provided through the Trautman Fund of The OSU Museum of Biological Diversity, The Ohio State University, and National Science Foundation DEB-1257796 to MD.

## Data Accessibility Statement

Raw sequence data, Python scripts, and all files for all analyses will be archived in Dryad upon final acceptance of this manuscript.

## Author Contributions

B.M.T. and M.D. conceived the study and collected samples, B.M.T. conducted laboratory work and analyzed the data. B.M.T. and M.D. wrote and edited the manuscript.

